# Group-level inference of information-based measures for the analyses of cognitive brain networks from neurophysiological data

**DOI:** 10.1101/2021.08.14.456339

**Authors:** Etienne Combrisson, Michele Allegra, Ruggero Basanisi, Robin A. A. Ince, Bruno Giordano, Julien Bastin, Andrea Brovelli

## Abstract

The reproducibility crisis in neuroimaging and in particular in the case of underpowered studies has introduced doubts on our ability to reproduce, replicate and generalize findings. As a response, we have seen the emergence of suggested guidelines and principles for neuroscientists known as *Good Scientific Practice* for conducting more reliable research. Still, every study remains almost unique in its combination of analytical and statistical approaches. While it is understandable considering the diversity of designs and brain data recording, it also represents a striking point against reproducibility. Here, we propose a non-parametric permutation-based statistical framework, primarily designed for neurophysiological data, in order to perform group-level inferences on non-negative measures of information encompassing metrics from information-theory, machine-learning or measures of distances. The framework supports both fixed- and random-effect models to adapt to inter-individuals and inter-sessions variability. Using numerical simulations, we compared the accuracy in ground-truth retrieving of both group models, such as test- and cluster-wise corrections for multiple comparisons. We then reproduced and extended existing results using both spatially uniform MEG and non-uniform intracranial neurophysiological data. We showed how the framework can be used to extract stereotypical task- and behavior-related effects across the population covering scales from the local level of brain regions, inter-areal functional connectivity to measures summarizing network properties. We also present an open-source Python toolbox called Frites^1^ that includes the proposed statistical pipeline using information-theoretic metrics such as single-trial functional connectivity estimations for the extraction of cognitive brain networks. Taken together, we believe that this framework deserves careful attention as its robustness and flexibility could be the starting point toward the uniformization of statistical approaches.

**Graphical abstract:** 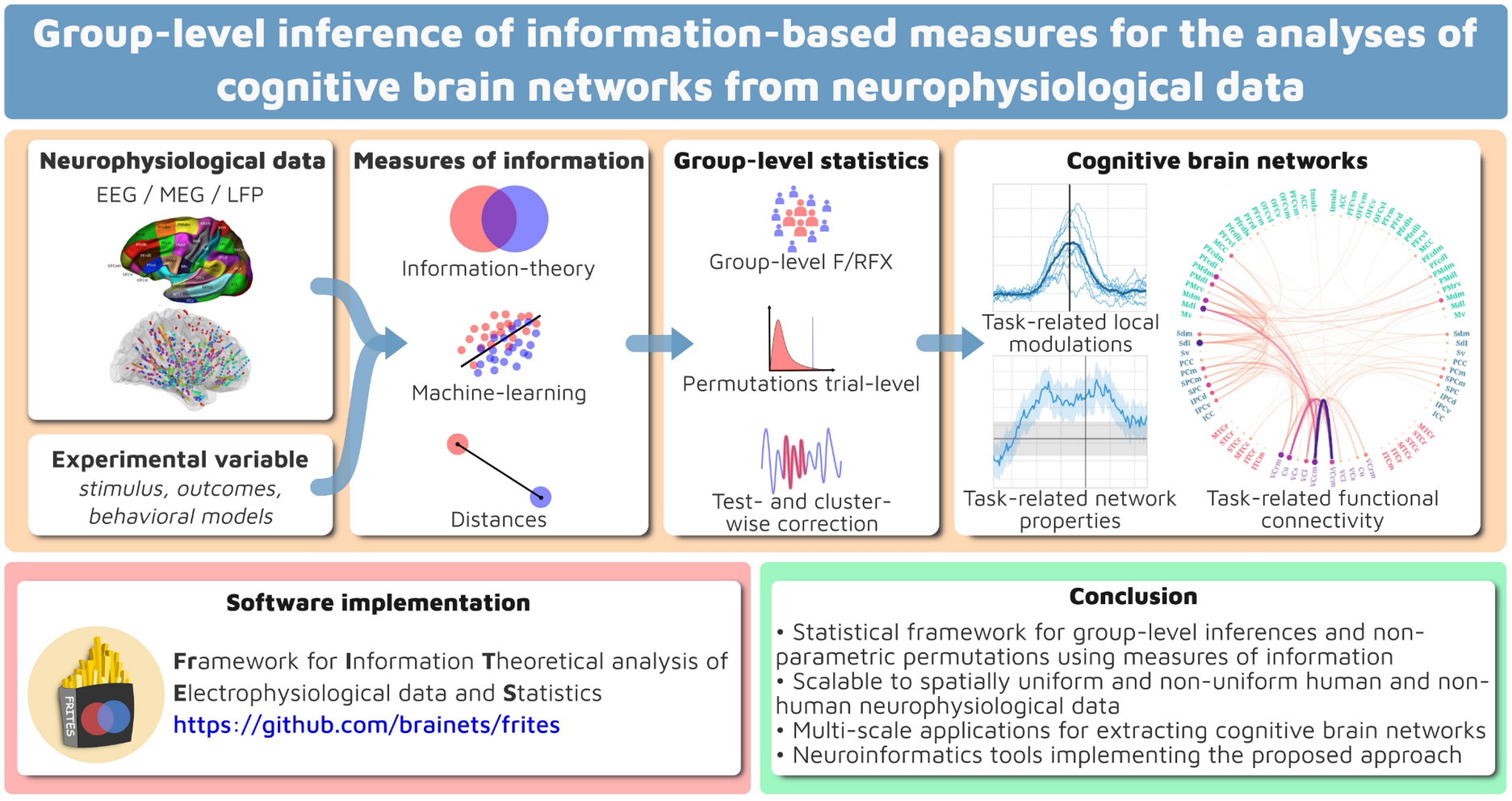

**Highlights:**

- Group-level statistics for extracting neurophysiological cognitive brain networks
- Combining non-parametric permutations with measures of information
- Fixed- and random-effect models, test- and cluster-wise corrections
- Multi-level inferences, from local regions to inter-areal functional connectivity
- A Python open-source toolbox called *Frites* includes the proposed statistical methods

## 1) Introduction

Modern theories suggest that cognitive functions emerge from the dynamic coordination of neural activity over large-scale and hierarchical networks (Bressler and Menon, 2010; Reid, 2019; Varela et al., 2001; von der Malsburg et al., 2010). A standard approach in cognitive brain networks analysis involves the characterisation of brain regions and inter-areal interactions that participate in the cognitive process under investigation (Battaglia and Brovelli, 2020; Bijsterbosch et al., 2020). Measuring the emergence of cognitive brain networks therefore underlies linking brain data to experimental variables, such as sensory stimuli or behavioral responses. Information-based measures can be used for this purpose as they are quantifying how much information is shared between brain data and experimental variables (Allefeld et al., 2016; Kriegeskorte et al., 2006; Kriegeskorte and Bandettini, 2007; Panzeri et al., 2017). Information theory (Ince et al., 2017; Panzeri et al., 2008; Timme and Lapish, 2018) and machine learning (Glaser et al., 2019) are two popular sets of measure of information. Information-based metrics are ideal tools to relate brain regions or network-level activity with task variables as they are returning quantities on a meaningful scale (like bits, percentage, variance explained etc.), they support multivariate data and they might not need to assume any specific type of dependency between variables.

The fMRI community has relatively standardized statistical pipelines to identify task-related functional activations at the population level. Some examples are established softwares like SPM (Friston et al., 1994) and analysis pipelines developed in international initiatives, such as the Human Connectome Project^1^ (Van Essen et al., 2013). While both also include pipelines on neurophysiological data, there is still less consensus on standard pipelines for investigating task-related neural representations and task-related network-level inference from neurophysiological data, leading to a larger heterogeneity in statistical group-level approaches. Such diversity mainly depends on a richer panel of modalities that can be used to record neurophysiological activity, each having unique spatiotemporal properties (e.g., EEG, MEG and intracranial EEG) and on differences in analysis goals (e.g., single-area versus network-level analyses). As an example, invasive intracranial recordings, which provide an exceptional signal-to-noise ratio at the single-trial level, are characterised by a spatially non-uniform sampling due to unique electrode implantations, which complexifies inferences at the population level and network level. On the other hand, M/EEG data provides an excellent technique for the analysis at the whole-brain level with the additional benefit of having an equal spatial sampling.

The reproducibility crisis in neuroimaging has many potential causes including analytic variability, statistical power and test-retest reliability (Poldrack et al., 2020). Taken together, these issues are limiting the ability of research groups to reproduce, replicate and generalize robust results (Arnold et al., 2019; Niso et al., *this issue*). Concerning neurophysiological M/EEG and intracranial data, several solutions have been proposed, from a uniformization of brain data organization through BIDS formatting (Holdgraf et al., 2019; Niso et al., 2018; Pernet et al., 2019) to overall good scientific practices and recommendations for data sharing, analysis and reporting results (Pernet et al., 2020, 2018). The problem of underpowered studies has been highlighted as one of the core limitation explaining the lack of reproducible results (Button et al., 2013; Ioannidis et al., 2014; Poldrack et al., 2017; Szucs and Ioannidis, 2017). In response, collaborative replication efforts like #EEGManyLabs (Pavlov et al., 2021) are attempting to replicate the findings of a subset of the most influential EEG studies using larger cohorts to improve confidence in most cited results. While such large-scale projects combined to data-sharing are currently tackling the low statistical power problem, the diversity of statistical approaches and multiple hypothesis testing have also been highlighted as one of the core limitations to reproducibility (Carp, 2012; Puoliväli et al., 2020). The flexibility of analysis procedures is one factor of increase of false positive results (Ioannidis et al. 2005; Gilmore et al., 2017). In other words, the larger the spectrum of methods the researcher can explore, the more likely he is to find one to validate an hypothesis. Therefore, there is a need for unifying statistical framework for extracting task-related neural representations both for single-area and network systems (Bassett et al., 2020) using information-based measures combined with a statistical layer accounting for the heterogeneity in the population (Mumford and Nichols, 2006) with the ability to adapt to the diversity of neurophysiological recordings.

Here, we address these issues by extending an existing statistical framework for group-level inferences on information-based measures (Giordano et al., 2017) to encompass neurophysiological recordings with both homogeneous spatial samplings across participants (or sessions), such as whole-brain source level data M/EEG data and spatially sparse recordings, such as intracranial sEEG, ECoG and LFPs. This framework supports any type of measure of information and can be used at the level of single brain area activity, connectivity-level inter-areal interactions (e.g., (un)directed functional connectivity measures) and network-level (e.g., on graph-theoretical measures). We tested the workflow on simulated data to compare fixed-versus random-effect group inference with test- or cluster-wise p-values correction. We then re-analyzed cortical high-gamma activity (HGA) data from different recording modalities: a MEG experiment (Brovelli et al., 2015), where we successfully retrieved the visuomotor network using information metrics from both IT and ML and an intracranial experiment (Gueguen et al., 2021), revealing the outcome-related activity of the anterior insula during probabilistic learning. Finally and as a contribution toward reproducibility, we also provide to the community the neuroinformatics tools wrapped in an open-source Python toolbox called Frites^2^ (*Framework of Information Theoretical analysis of Electrophysiological data and Statistics*). This package includes the proposed group-level statistical framework such as functions for the estimation of functional connectivity. By default, Frites is using metrics from the IT but it can be extended to any type of measure of information.

## 2) Methods

### 2.1 Statistical framework for group-level inferences of information-based measures

The main objective of the current work was to develop a non-parametric framework to assess the group-level statistical significance on measures of information computed on neural correlates of individual brain areas or interareal relations, such as functional connectivity (FC) measures. This framework concerns task-designs with independent repeated measurements (i.e. trials or epochs). It allows investigating condition-related (i.e. stimulus or outcomes) effects (i.e. information shared between continuous brain data and a vector of discrete integers specifying the category to which each trial belongs). It also allows investigating behavior-related effects (i.e. information shared between continuous brain data with continuous behavioral models or psychometric measurements). In the rest of the manuscript, we will use the term “*task-related*” to encompass both condition- and behavior-related effects. The proposed framework (**Figure 1**) relies on two main steps : *(i)* the estimation of the true and permuted effect size using a measure of information between neural data *X* and an experimental variable *Y*; *(ii)* statistical inference at the test-wise (i.e., at a single time-point) or cluster-level (Maris and Oostenveld, 2007).

**Figure 1.**
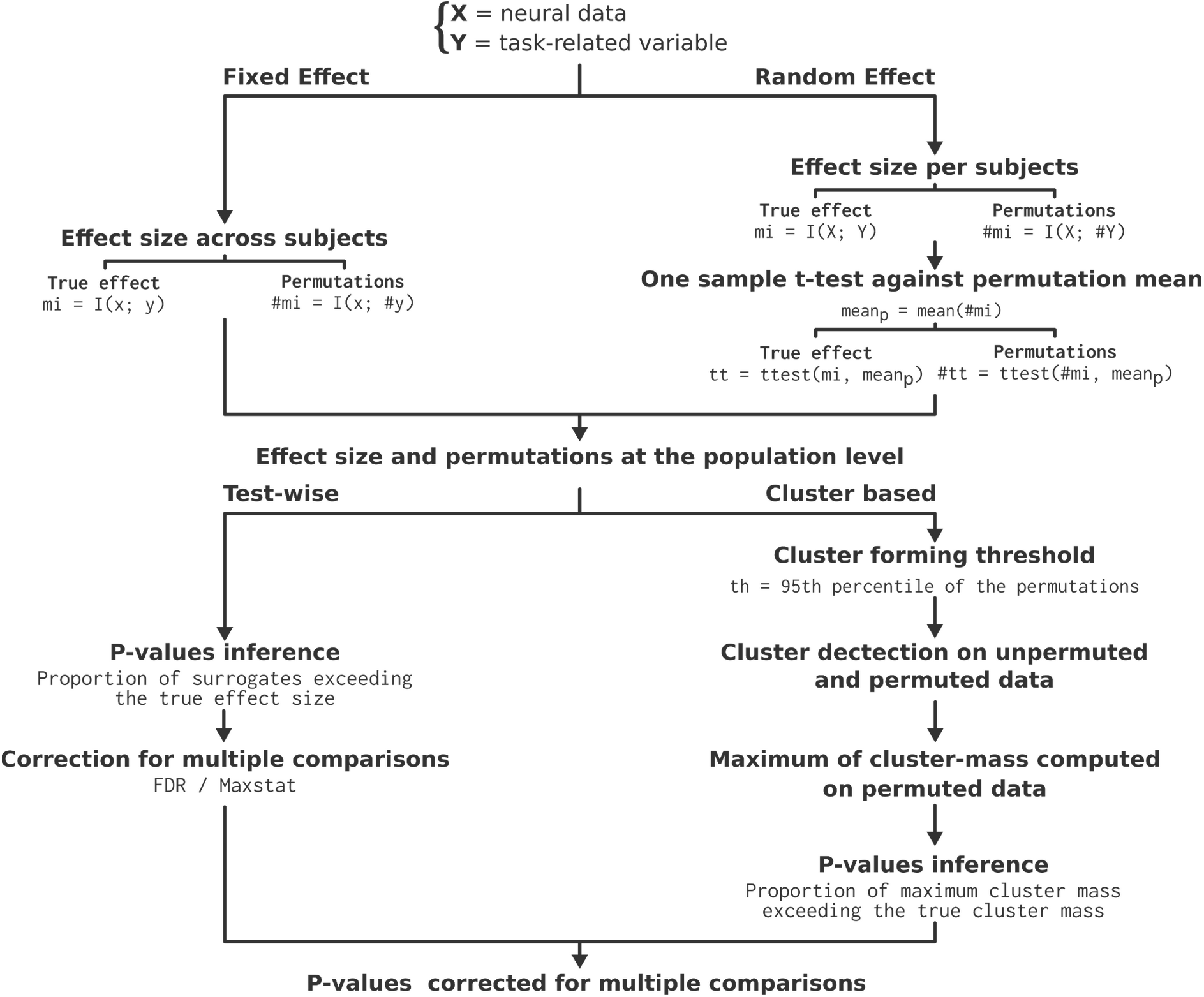
Statistical framework for group-level inferences using information-theoretic measures (I=Mutual-Information) to extract task-related neural activity or functional connectivity. The upper branches describe the fixed and random effect implementations, both leading to a measure of a true effect size and the estimation of the null distribution. Then, starting from the effect size and the permutations, bottom branches describe how both are used to infer and the p-values corrected for multiple comparisons using either test- or cluster-wise corrections.

#### 2.1.1. Effect size estimation and permutation tests for fixed and random effect models

The estimation of effect size on a population depends on assumptions about the underlying distribution and inter-subject variability. A fixed-effect (FFX) is a single-level model assuming that each subject brings the same fixed contribution to the observed effect (Friston et al., 1999). It extracts the typical characteristics of a sample, which explain why it is massively used for reporting case studies. A FFX at the population level can be estimated by computing the effect size across all concatenated subjects and trials (Penny and Holmes, 2007). This is equivalent to considering the data as coming from a single subject, and discards random variations across subjects. The true effect size for FFX is therefore computed on the concatenated trials across participants using a measure of information (see top panel of **Figure 1**, left branch). Significance for fixed- or random-effects was performed using a nonparametric approach based on permutations. Since the aim is to quantify task-related effects, permutations are performed by randomly swapping the task-related variable, as usually performed in other fields (Combrisson and Jerbi, 2015). Permutation-based significance testing shortcuts distributional assumptions by relying on the *exchangeability* hypothesis i.e. that the observations are exchangeable under the null hypothesis (Nichols and Holmes, 2002). In our approach, the null hypothesis is that no statistical relationship and significance exists between the trial-by-trial modulation in brain activity *X* and task-related variable *Y*, as measured by means of information theoretical or ML metrics. For a permutation test, that means that if the null hypothesis is true, the labels assigning task-variables to trials are interchangeable. For FFX analyses, the estimation of the null distribution of effect sizes assumes exchangeability of results from different participants and trials, and it is implemented by means of random permutation of data across both. Thus, for FFX, the null effect size corresponds to the information between brain activity *X* and randomly-permuted task-related variable *Y*.

While FFX shines by its simplicity, conclusions that can be drawn only concern the subjects that constitute the available sample as a whole, and therefore can not be generalized to the population-level. On the contrary, a random-effect (RFX) is a hierarchical model consisting in : *(i)* extraction of single-subject effect size and *(ii)* fitting a model at the population level. Here, the sample of subjects is considered as a subsample of a broader population and then, if new subjects are added, their effect size should fall into the estimated model (see top panel of **Figure 1**, right branch). First, the information shared between the neural data *X* and the experimental variable *Y* is computed at the single-subject level. For the permutations, we assume exchangeability of results from different trials only, and permute trial-specific results independently for each participant. To then form the true effect size and the null distribution at the population level, we used a one-sample t-test, computed across single-participants, against the null population mean (mean_p in **Figure 1**). The null population mean was computed by taking the average of the trial permuted measures of information across participants. This produced true and permuted t-values that quantified the true and null effects size for the RFX analysis, respectively. As information-based metrics are often subject to bias, it is not correct to perform a parametric population level t-test against a specific value (such as 0). Obtaining a non-parametric population null distribution from within-participant trial permutations avoids this problem, as the permutation values are obtained from the same number of samples, and so will be subject to the same level of bias.

In the end, both FFX and RFX procedures led to the true effect size and the distributions of permutations corresponding to the effect size that can be obtained by chance. However, for the FFX, the summary statistics at the population level is expressed in the unit of the information-based method (i.e in bits when using IT metrics or, for example, in percentage for ML decoding) while for the RFX it corresponds to T-values (**Figure 1**).

#### 2.1.2. Inferring p-values and correcting for multiple comparisons

In order to assess whether the estimated effect size significantly differs from the chance distribution and to correct for multiple comparisons (Cao and Zhang, 2014; Lindquist and Mejia, 2015; Nichols and Hayasaka, 2003), we implemented both test- and cluster-wise statistics (Maris and Oostenveld, 2007). In the case of test-wise correction, p-values are defined as the proportion of surrogates that are exceeding the observed true effect size estimation. The p-values are then corrected for multiple comparisons using the False Discovery Rate (Genovese et al., 2002) or the maximum statistics, which provides a threshold that controls family-wise error rate (Holmes et al., 1996; Nichols and Holmes, 2002). For cluster-level statistics, the cluster forming threshold is defined as the 95th percentile across all of the permutations. Such threshold is used to identify the clusters on both the true and permuted data. In order to infer the p-values and to correct for multiple comparisons, a null distribution composed of the maximum of the cluster mass is built and the corrected p-values are obtained by assessing the proportion of cluster masses computed on the surrogates that exceed the cluster mass of the true effect size.

For simplicity, the description above about the statistical framework is considering the participant as the fixed or random variables. The same description still holds if sessions are used instead or, in the context of intracranial data, the recording contacts across participants (Gueguen et al., 2021).

### 2.2 Measuring information for the analysis of cognitive brain networks

Information theory (Cover and Thomas, 1991) represents one of the most successful and promising data-driven frameworks to quantify task-related information in neural signals (Borst and Theunissen, 1999; Quiroga and Panzeri, 2009) and it has been extensively applied to neurophysiological signals (Cogan et al., 2011; Giordano et al., 2017; Gross et al., 2013; Ince et al., 2016; Jeong et al., 2001; Magri et al., 2009; Schyns et al., 2011) and neuroimaging studies (Ince et al., 2017; Ostwald and Bagshaw, 2011; Panzeri et al., 2008; Timme and Lapish, 2018; Wibral et al., 2015). In addition, information theoretic measures based on the Wiener-Granger principle (Bressler and Seth, 2011; Brovelli et al., 2004; Ding et al., 2006; Seth et al., 2015), such as Transfer Entropy (Schreiber, 2000) and Directed Information (Massey, 1990), provide means to quantify network-level directional influences and can capture arbitrary (linear and nonlinear) time-lagged conditional dependencies between neural signals (Besserve et al., 2015; Ince et al., 2015; Vicente et al., 2011). Although our statistical approach can be applied to all information-based metrics, we focused our analysis and application to information theoretical quantities, such as mutual information (MI) and conditional MI (CMI) measures (Ince et al., 2012). Note that the permutation strategy for CMI estimates (i.e. I(X; Y|Z)) also consists in randomly permuting the *Y* variable across trials and leaving both *X* and *Z* untouched.

The standard method to compute entropies for the calculation of mutual information (MI) and conditional MI (CMI) is the binning method. Such “direct” method consists in binning each univariate variable and collecting the bin counts, so to approximate the joint probability distribution *P*(*X,Y*) by the multidimensional histogram (Beirlant et al., 1997; Treves and Panzeri, 1995). However, binning strongly depends on the number of samples and suffers from the curse of dimensionality (Geman et al., 1992). Binless strategies have also been proposed, such as Kernel Density Estimation (KDE) methods (Moon et al., 1995). However, KDE estimations suffer the same limitations of direct methods linked to the curse of dimensionality (Gramacki, 2018; Scott, 2015) typical of high-dimensional datasets, such as neurophysiological data. A promising alternative based on semi-parametric estimation techniques has been recently proposed, namely the Gaussian Copula Mutual Information (GCMI) (Ince et al., 2017). In short, MI does not depend on the marginal distributions of the variables, but only on the copula function which describes their dependence. GCMI therefore first requires transforming the inputs so that the marginal distributions are a standard normal. This copula-normalisation step involves calculating the inverse standard normal cumulative density function (CDF) value of the empirical CDF value of each sample, separately for each input dimension, before using a parametric, bias-corrected, Gaussian MI estimator (i.e., sumrank computation). The included parametric bias-correction is an analytic correction to compensate for the bias due to the estimation of the covariance matrix from limited data (i.e. here, limited number of repetitions or trials). Since this parametric correction only depends on the number of trials, the same value is going to be used for both permuted and nonpermuted data. Therefore, this bias correction only impacts the estimated effect size but has no effects on statistical results. The GCMI provides a lower bound estimate of the MI, without making any assumption on the marginal distributions of the input variables (Ince et al. 2017; Ma and Sun 2011). It is important to stress that the rank-based copula-normalisation preserves the relationship between variables as long as this relation is monotonic (i.e., strictly increasing or decreasing). For example, if the Y variable is a parabolic function of X, the ranking is not preserved, and thus the relation between them can not be detected by the GCMI. In other words, for 1D variables the GCMI is capable of detecting linear and nonlinear relations, as long as the relation is monotonic.

Nevertheless, GCMI has several advantages for brain network analysis. The first advantage is the simplicity of the computation. This renders the algorithms applicable to large datasets with hundreds of variables (e.g., brain regions) and it makes it suitable for computationally-intensive nonparametric permutation tests. The second advantage is the ability to estimate entropies on few data samples. This property allows the estimation of MI and CMI from short time series containing hundreds of time points, e.g., single trials or across trials. A third advantage is the possibility to compute MI and CMI between a combination of discrete and continuous variables, that can either be univariate or multivariate, with resulting effect sizes on a common and interpretable scale. This provides a tight link between information theoretic quantities and classical statistical approaches used in neuroscience. For example, if both variables are continuous and univariate (e.g., two timeseries, one in the parietal and one in the occipital), MI is related to the generalized Pearson correlation. If *X* is a continuous variable (e.g., the high-gamma activity in the visual cortex) and *Y* is discrete and univariate (e.g., task-related variable), their MI is equivalent to a Student t-test and it is related to a decoding machine-learning analysis where *X* is the array of samples and *Y* the vector of labels. Furthermore, the CMI produces an equivalent of partial correlation. For a more detailed description of the relation between information theoretic quantities and other statistical approaches, see the Table I in Ince et al. 2017. It should be stressed out that in the case of MI between a continuous and a discrete variable containing more than two categories (e.g. four visual stimuli like cat, dog, penguin and horse pictures) only a subsample of stimuli (e.g. cat and horse) can drive the MI to higher values. To disentangle the contribution of each stimulus, decoding analysis and inspection of the confusion matrix could be used in place of the MI.

### 2.3 Software implementation

We implemented the framework presented in this paper in an open-source python package called *Frites*^3^ (*Framework for Information Theoretical analysis of Electrophysiological data and Statistics*). The key objective of *Frites* is to identify task-related neural activity or cognitive brain networks through information theory and also to be able to draw inferences at the group level. *Frites* contains two important classes. The first one is the workflow of MI (frites.workflow.WfMi^4^) to quantify the statistical dependency between the neural data and a task-related variable. The *WfMi* can be used to investigate the quantity of information shared between continuous and discrete variables (e.g. brain data and two or more experimental conditions or stimulus types), between two continuous variables (e.g. brain data and with behavioral-models or psychometric measurements) or between two continuous variables conditioned by a third discrete one (conditional mutual information). This workflow estimates the effect-size and the surrogates as shown in the top part of **Figure 1**. By default and for efficient computations, *Frites* is using a tensor-based implementation of the GCMI. However, custom estimators can be defined and provided using the frites.estimator.CustomEstimator object to estimate effect sizes with for example decoders, measures of distance, correlations etc. Then, the effect-size and the surrogates are plugged into the second important workflow for group-level statistical inferences (frites.workflow.WfStats^5^), which corresponds to the bottom part of **Figure 1**. Cluster-based correction is performed using the open-source software MNE-Python (Gramfort et al., 2013). One important aspect is that, in contrast to fMRI data, electrophysiological recordings do not necessarily have a spatial contiguity (e.g sparse intracranial data). Therefore, when using cluster-based corrections, the cluster entry threshold is inferred across all time and space bins but then, the clusters are only detected across time. Finally, for correcting for multiple comparisons, the maximum of the cluster masses is taken across time and space.

### 2.4 Numerical simulations

We performed numerical simulations to generate artificial task-related data and then measure the performance of the group-level statistics when combined with information-based measures. More precisely, we tested how FFX and RFX combined to test- or cluster-wise corrections for multiple comparisons are affected by various parameters. The simulated data consisted in two variables: a 3D multivariate variable *X*, which simulates the spatio-temporal structure of electrophysiological data with the dimensions describing the number of trials, the number of brain regions and the number of time points and a univariate variable *Y*, reflecting a task-related continuous variable with the same number of trials as in *X*. Both *X* and *Y* were defined as standard random Gaussian variables without temporal autocorrelation. Then, for a given brain region *b*, we introduced a statistical dependency between those two variables during a temporal interval [*t*_1_*;t*_2_] :

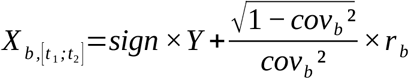

where *r* is a random gaussian noise of unit variance, *sign* that can either be −1 or 1 and *cov* a correlation parameter with values between 0<*cov*≤1 for controlling the correlation strength between *X* and *Y*. If *cov* is 1 a perfect correlation is observed between *X* and *Y*. On the contrary, the closer to zero *cov* is, the less the variables are correlated. As a result, the correlation between both variables exist only during a certain temporal period and can also vary across brain regions. However, the direction of the correlation (i.e. correlation or anticorrelation) was the same across all brain regions.

We defined three scenarios associated with different task-related effects whose amplitude and duration varied across time and space (**Figure 2**). For each scenario, a boolean ground-truth array was built with 0 and 1 indicating the absence or presence of effect, respectively, at a specific time and in a specific region. The objective was then to compare such a ground-truth array with the output statistics.

**Figure 2.**
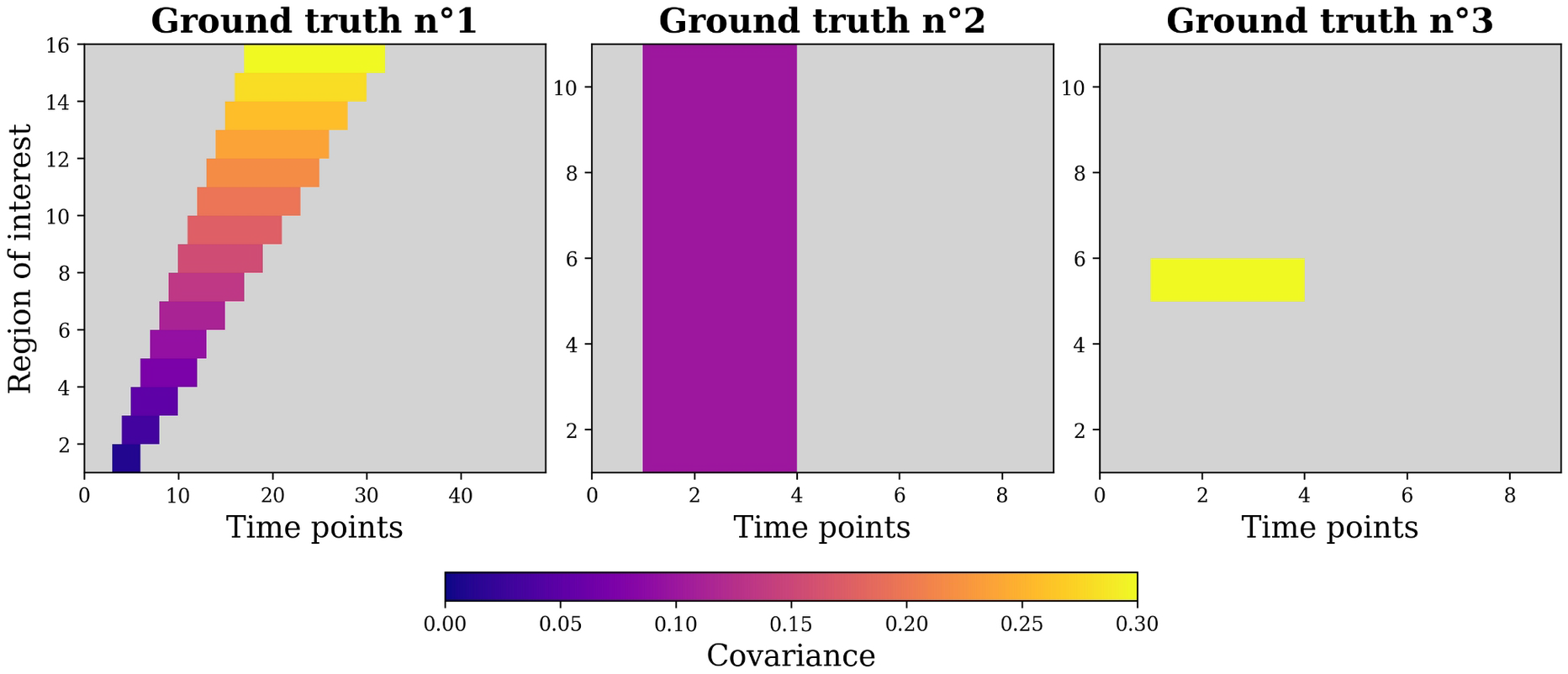
simulated spatio-temporal ground truths (GT) reflecting the presence and the strength of the statistical dependency between the data and the stimulus. The simulated effect is distributed across space (region of interest) and time points with a varying color-coded covariance. The higher the covariance, the easier the effect should be to be retrieved. The first GT simulates a spatially broad effect, with varying covariances and temporal cluster lengths. The second GT simulates a weak and diffuse effect and the third, a strong and focal effect. Gray parts symbolize the absence of effect.

In addition to the above ground-truths describing various spatio-temporal profiles, we also measured the impact of four intertwined parameters of the simulated data, namely *(i)* the size of the population (i.e the number of subjects) *(ii)* the within-subject size (i.e the number of trials per subject (Baker et al., 2020)), *(iii)* the proportion of subjects having the effect and *(iv)* the inter-subject consistency or ISC. The ISC varied between 50 and 100% and characterized the proportion of subjects with either positive (*sign*=1) or negative (*sign*=−1) correlation between the data and the stimulus. As an example, let us consider a dataset composed of 10 subjects, among which 80% has the effect (8 subjects out of 10). In such a case, an ISC of 50% means that 4 subjects have positive correlations, 4 have negative correlations and 2 have no effect. An ISC of 100% means that, among the subjects having the effect, all of them have positive correlations.

In its most general definition, mutual information is capable of capturing any type of relation. However, Gaussian copula MI between two continuous variables makes the assumption of a monotonic relation between the variables. When using FFX group-level approach, the MI is computed on data concatenated across subjects : this means that when positive and negative correlation coexist at the group level, the monotonicity assumption is broken. This does not occur in the RFX, as the latter computed MI independently for each subject. As a consequence, a low ISC (i.e with a balanced proportion of subjects having positive and negative correlations) should impact the FFX more than the RFX.

The MI was computed across trials, at each time-point and brain region. The p-values estimated at the group-level were inferred using the above-described permutation approach. We considered a significance threshold of p<0.05 to compute the fraction of true and false positives, and true and false negatives (TP, FP, TN, and FN, respectively). Those fractions were inferred by comparing the detected significant effects with the ground-truth. As an example, for the RFX model with FDR correction, we used the non-parametric framework to find significant effects at p<0.05. This step leads to a spatio-temporal boolean matrix, with the same shape as the ground-truth, where 1 and 0 respectively refer to significant nonsignificant effects. Then the fraction of FP corresponds to the spatio-temporal overlap between effects detected as significants (i.e. ones in the boolean matrix) and the presence of a true effect in the simulated ground-truth. Similarly, TN fractions correspond to the overlap between zeros in the boolean matrix and the absence of effect in the ground-truth. This procedure was repeated ten times in order to decrease the variability. We computed the *sensitivity* (or true positive rate) and *specificity* (or true negative rate) of the various approaches to group-level inference (Nichols and Hayasaka, 2003; Shapiro, 1999) defined as:

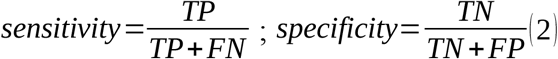

In practice, *sensitivity* measures the overall accuracy for the detection of a true effect, and *specificity* measures the overall accuracy for the detection of the true absence of an effect. We then pooled information across measures of *sensitivity* and *specificity* to compute a summary measure of performance for the group-level method. To this purpose, we considered the Matthews Correlation Coefficient (MCC) (Matthews, 1975) defined as :

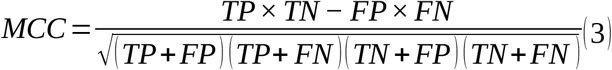

where MCC can assume any value in the [−1, 1] range, with 1 measuring a perfect performance, −1 measuring a perfect negative correlation and 0 a chance-level performance (Chicco and Jurman, 2020; Mensen and Khatami, 2013; Roels et al., 2016; Vihinen, 2012). The MCC reported in **Figure 3** and **Figure 4** was inferred by computing the fractions of T/FP and T/FN across the three ground-truths, across all time and space bins and across the ten repetitions. An example illustrating the simulated ground-truths, the computations of the statistics at the group-level such as the definition of the fractions and statistical metrics can be found in the documentation of the software^6^.

**Figure 3.**
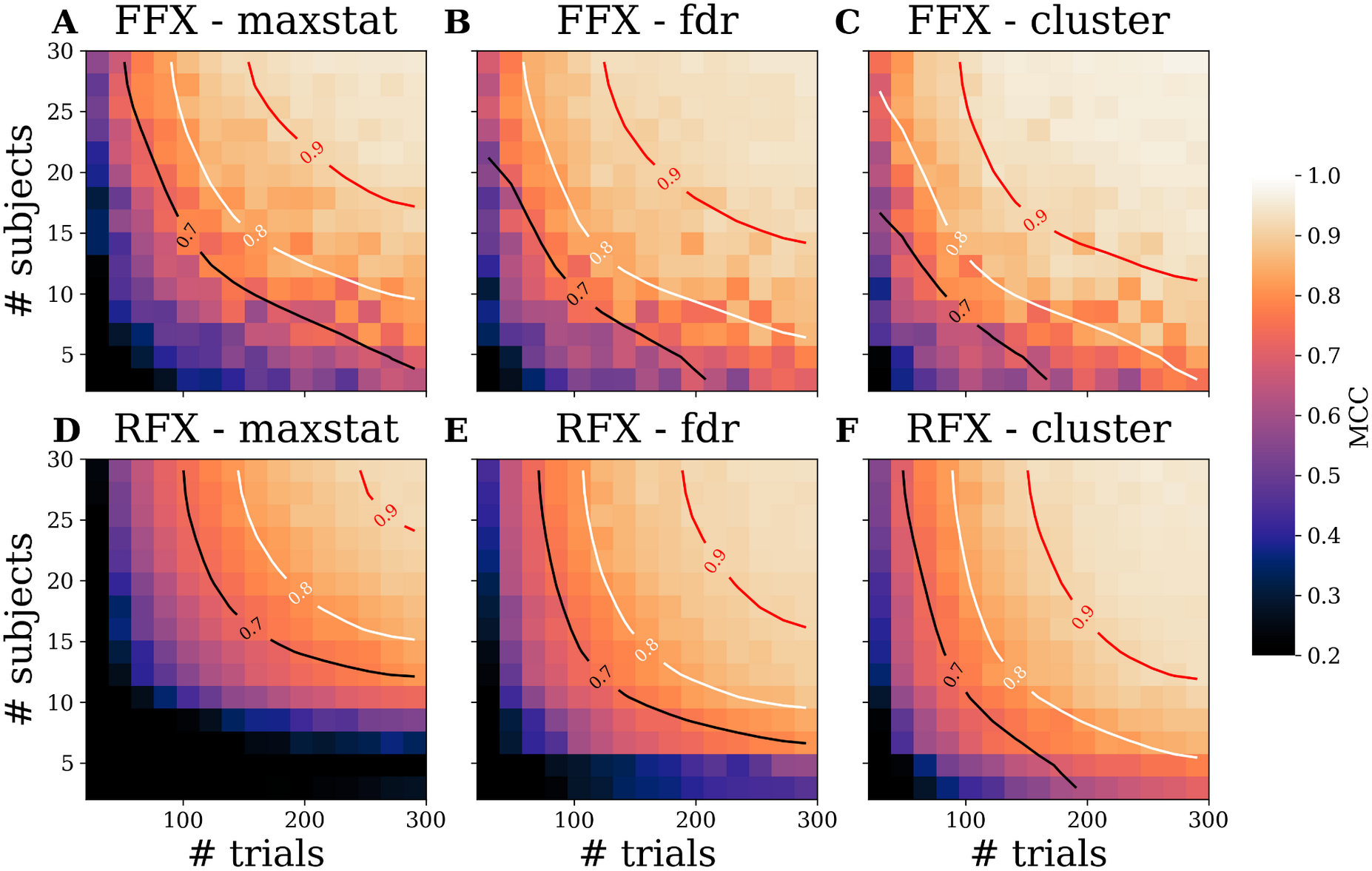
Comparison of group-level approaches on simulated data with varying number of trials (from 20 to 300) and number of subjects (from 2 to 30). The comparison includes the FFX (*A-C*) and the RFX (*D-F*), each combined with the three methods for correcting the p-values for multiple comparisons (maximum statistics, FDR and cluster-based) across time and brain regions. The MCC is computed across all three ground truths and the 10 repetitions, with 80% of the subjects having the effect and with a fixed ISC of 90%. Black, white and red lines respectively represent a MCC of 0.7, 0.8 and 0.9.

**Figure 4.**
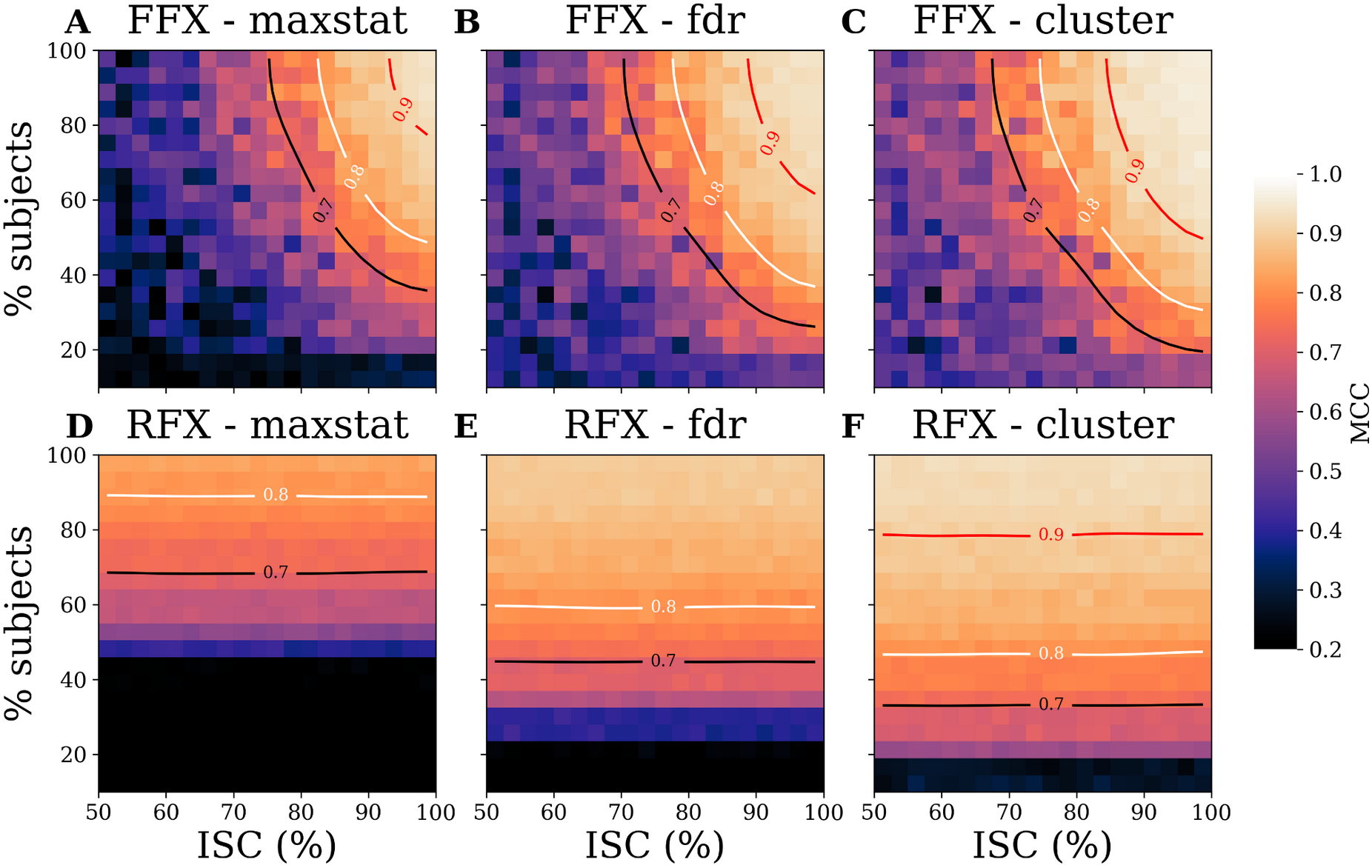
Comparison of group-level approaches on simulated data with varying inter-subject consistency (from 50 to 100%) and proportion of subjects having the effect (from 10 to 100%). The comparison includes the FFX (*A-C*) and the RFX (*D-F*), each combined with the three methods for correcting the p-values for multiple comparisons (maximum statistics, FDR and cluster-based) across time and brain regions. The MCC is computed across all three ground truths and the 10 repetitions, across 15 subjects and 200 trials. Black, white and red lines respectively represent a MCC of 0.7, 0.8 and 0.9.

### 2.5 Validation on MEG and intracranial sEEG electrophysiological data

To validate the proposed statistical approach, we reanalysed human brain data previously published using standard parametric approaches. We analysed: i) source-level high-gamma activity (HGA) estimated from MEG data recorded during the execution of a visuomotor-mapping learning task (Brovelli et al., 2017, 2015) and (ii) HGA estimated from intracranial recordings (stereotactic electroencephalography, sEEG) acquired during a probabilistic learning task (Gueguen et al., 2021).

#### 2.5.1. MEG high-gamma activity during a visuomotor-mapping learning task

We reanalysed HGA (60-120Hz) estimated at the source level on ten right-handed healthy subjects performing a visuomotor-mapping task (Brovelli et al., 2015) consisting in executing finger movements instructed by a visual stimulus (digit 1 instructed the execution of the thumb, digit 2 for the index finger, digit 3 for the middle finger, and so on). Each participant performed two sessions of 60 trials each (total of 120 trials). MEG recordings were performed using a 248 magnetometers system (4D Neuroimaging, magnes 3600). Single-subject cortical parcellation was performed using the *MarsAtlas* brain scheme (Auzias et al., 2016). Single-trial and z-scored (with respect to a baseline period from −0.5 to −0.1 s before stimulus onset) HGA time-series aligned on finger movement were estimated for each *MarsAtlas*’ parcel, as described in previous studies (Brovelli et al., 2017). The goal of the original publication was to isolate the brain regions involved in the performance of arbitrary visuomotor associations. To this end, authors used linear mixed-effect models to contrast the HGA activity occurring during baseline against the HGA while performing the visuomotor association. The dynamic functional connectivity (DFC) was computed using the Pearson correlation over sliding windows of 500 ms stepped every 10 ms.

In the present study, we investigated whether it was possible to retrieve the visuomotor-related cortical brain regions and network using only information-based measures combined with the proposed group-level statistical framework. To this end, we computed the MI between the single-trial HGAs and a discrete label vector describing whether the trials belonged to one or the other separated periods (i.e. the baseline (−0.6 to −0.1s) or visuomotor-related periods). The permutation scheme consisted in randomly shuffling this label vector. To highlight the generalizability of the proposed framework to other measures of information, we also used machine-learning (ML) outputs. For the ML analysis, we used a Linear Discriminant Analysis classifier (LDA) with a stratified 10-fold cross-validation to split the data into training and testing sets. The LDA was used to decode whether the HGA was coming from the baseline or visuomotor-related period. A new classifier was systematically defined at each time point to produce time-resolved analysis. For summarizing the quality of predictions, we used the area under the curve (AUC) metric. ML analysis was performed using the Python software scikit-learn (Pedregosa et al., 2011).

For network-level analyses, we computed the single-trial DFC on HGA using the MI during the 500ms baseline period and using sliding windows of 500 ms stepped every 10 ms during the visuomotor-related period. We then computed the MI and p-values between the DFC estimated during the 500ms baseline and visuomotor-related periods using a RFX approach with FDR correction. To illustrate how the choice of the group-level approach is impacting the number of significant pairwise links, we performed the same contrast of DFC during baseline vs. DFC during visuomotor related period using all possible combinations between F/RFX models and correction for multiple comparisons. Finally, to illustrate the possibility of the proposed framework to analyse graph-theoretical metrics computed on DFC matrices, we investigated task-related differences of frequently found network measures (Bullmore and Sporns, 2009) by means of network global efficiency, modularity and assortativity using the Brain Connectivity Toolbox (Rubinov and Sporns, 2010). To this end, we computed each measure at the single-trial level on the binarized FC matrix estimated during the baseline and on each binarized FC matrix estimated during the visuomotor-related period using the sliding windows. Since it is not the core point of this paper, the binarization of the DFC matrices was performed using an arbitrary threshold to keep 50% remaining links. A more robust analysis would require investigating multiple thresholds (Rubinov and Sporns, 2010). As a result, we got on one side a single vector reflecting for example the network efficiency of each trial during the baseline period and on the other side, the 2D matrix of single-trial and dynamic network efficiency estimated during the visuomotor-related period. Finally, we computed the mutual-information between each graph metric and a discrete vector specifying whether each trial belongs to the baseline of visuomotor period using a RFX model with cluster-based correction.

#### 2.5.2. Intracranial sEEG recordings during a probabilistic learning task

We then investigated outcome-related effects in intracranial HGA, previously observed using general linear models (GLM). Previous analyses revealed differently modulated activity during the reward and punishment conditions in the anterior insula (aINS) (Gueguen et al., 2021). The dataset was composed of twenty patients with medically-intractable epilepsy, implanted with five to seventeen stereotactic electroencephalography (sEEG) electrodes. Patients performed a probabilistic instrumental learning task adapted from previous studies (Palminteri et al., 2012; Pessiglione et al., 2006). The task consisted in the presentation of visual stimuli leading either to monetary gains or losses. Subjects were instructed to maximize their financial payoff, while considering reward-seeking and punishment-avoidance strategies as equally important. During the task, four pairs of visual abstract cues taken from the Agathodaimon alphabet were presented on a screen. The four cue pairs were divided in two conditions (2 pairs of reward and 2 pairs of punishment cues), associated with different pairs of outcomes (winning 1€ versus nothing or losing 1€ versus nothing). Subjects performed three to six sessions during which each pair was presented 24 times, leading to a total number of trials across sessions between 288 and 576. SEEG data preprocessing was conducted according to routine procedures described in previous studies (Bastin et al., 2016; Combrisson et al., 2017; Jerbi et al., 2013, 2009). These included signal bipolarization, where each electrode site was re-referenced to its direct neighbor. Bipolar rereferencing can increase sensitivity and reduce artefacts by canceling out distant signals that are picked up by adjacent electrode contacts (e.g., mains power). Next, electrodes contaminated with pathological epileptic activity were systematically removed using visual inspection and time-frequency decompositions. Time-frequency analyses were carried out using MNE-Python software (Gramfort et al., 2013). Single-trials HGA aligned on outcome presentation were estimated using a DPSS tapers (central frequency of 100hz, 15 cycles, time bandwidth of 15 leading to a frequency range of [50, 150]Hz). A more detailed description of the data acquisition and the task design have already been described elsewhere (Gueguen et al., 2021). In the present study, we focused on the anterior insula and investigated whether the HGA was differently modulated during the reward and punishment conditions. To this end, we conducted two separate analysis were we first computed the amount of information shared between the HGA and the discret vector of outcomes during the rewarding condition (i.e. with the outcomes +0€ and +1€) and then during the punishment condition (i.e. with the outcomes −1€ and −0€). For the group-level statistics, we used the same approach as the original publication, a RFX model across contacts (i.e. within a brain region, merge the contacts coming from multiple subjects, compute the MI per contact and then perform the t-test across the contacts).

## 3) Results

In the following sections, we will first present the results of numerical simulations assessing the performance of the workflows in statistically detecting various spatio-temporal profiles. We will then present the results obtained from the reanalysis of cortical HGA estimated from MEG data during visuomotor mapping and from intracranial data during reinforcement learning.

### 3.1 Numerical simulations: accuracy comparison of the group-level approaches

Three ground-truth scenarios (**Figure 2**) were simulated to compare the two group-level approaches (FFX and RFX) combined with one of the three methods for correcting p-values for multiple comparisons across time and brain regions, namely, the maximum statistics, the FDR or the cluster-based, respectively. We also varied the number of subjects and trials (**Figure 3**), such as the proportion of the subjects having the effect and the intersubject consistency or ISC (**Figure 4**).

**Figure 3** shows the Matthews Correlation Coefficient (MCC), an overall measure of performance (Sec. 2.4), as a function of the number of simulated subjects and trials. Overall, the FFX (**Fig. 3A-C**) leads to slightly higher values of MCC compared to the RFX (**Fig. 3E-F**) and across the three correction methods. This translates into a decrease in the data size requirements. For example, using the maximum-statistics at 200 trials, the FFX (**Fig. 3A**) reaches a MCC of 0.8 using approximately 12 subjects while the RFX (**Fig. 3D**) requires around 20 subjects. Additionally, for both the FFX and RFX, the cluster-based correction (**Fig. 3C and F**) performed systematically better than maximum statistics (**Fig. 3A and D**) and FDR (**Fig. 3B and E**). Note that we fixed the percentage of subjects having the effect at 80% and the ISC at 90%. Therefore, this set of parameters simulates a scenario where the relation between the data and the stimulus is relatively uniform across subjects, which is in favour of the FFX’s assumptions.

We then investigated the dependence on an additional parameter relevant in group-level analyses, namely the heterogeneity across subjects, which was simulated by varying the inter-subject consistency (ISC). **Figure 4** shows the MCC values as a function of the ISC (varying from 50 to 100%) and the percentage of subjects having the effect that varies (varying from 10 to 100%). A clear difference in dependence between the FFX and the RFX was observed as a function of the ISC. The MCC under the RFX model was stable for varying values of ISC (**Fig. 4D-F**). On the other hand, the FFX required that at least 80% of the subjects have the same type of effect in order to reach a MCC of 0.8 (i.e 80% with positive or negative correlations between the simulated data and the stimulus) (**Fig. 4A-C**). Nevertheless, for ISC values approximately higher than 90%, the MCC scores were larger in the FFX than in the RFX setting. The second main difference concerned the correction for multiple comparisons. **Fig. 4C and F** show that cluster-based correction outperformed the maximum statistics and the FDR. Interestingly, the correction had a strong impact on the required proportion of subjects having the effect for the RFX (larger than 80% with the maximum statistics and ~50% for the FDR and cluster-based).

### 3.2 Brain data: extracting task-related cognitive brain networks using information-based measures from human neurophysiological data

To validate the proposed statistical approach on human neurophysiological data, we reanalysed two previously-published datasets, which were analysed using standard parametric approaches. We analysed: *(i)* source-level high-gamma activity (HGA) estimated from MEG data recorded during the execution of visuomotor-mapping task (Brovelli et al., 2017, 2015) and *(ii)* high-gamma activity (HGA) computed from intracranial recordings (stereotactic electroencephalography, sEEG) acquired during a probabilistic learning task (Gueguen et al., 2021). For both modalities, we used the single-trial high-gamma activity (HGA) as the neural marker modulated by the task.

#### 3.2.1. Visuomotor-related brain network and MEG high-gamma activity

We computed the MI between the single-trial HGAs and a discrete vector of conditions (baseline versus visuomotor-related periods). The aim was to identify which brain areas displayed visuomotor-related changes in HGA. We then performed a random-effect group-level analysis, where the mutual information was estimated for each subject and then a t-test was applied across subjects. The p-values were corrected for multiple comparisons using a cluster-based approach.

We reproduced the main findings of a previous study (Figure 2 in Brovelli et al., 2017) revealing a visuomotor-related large-scale network **(Figure 5)** involving bilateral dorsolateral and dorsomedial motor regions (Mdl, Mdm), premotor (PMdl, PMdm) and somatosensory areas (Sdl, Sdm). Parietal cortices also displayed differences in both hemispheres in the medial and superior parietal cortex (PCm, SPC, SPCm) such as in the mid, posterior and inferior portions of the cingulate cortex (MCC, PCC, ICC). Some activations were also significant only in the left hemisphere, namely the caudal dorsomedial part of the prefrontal cortex (PFcdm), the ventral somatosensory (Sv) such as the dorsal and ventral parts of the inferior parietal cortex (IPCd, IPCv). Finally, we found large differences in HGA within the occipital lobe, especially in the rostral medial and caudal medial part of the visual cortex (VCcm, VCrm) and cuneus (Cu) of both hemispheres. To a lesser degree, the superior and lateral parts of the visual cortex (VCs, VCl) also exhibited differences (Figure 5). It should be noticed that, around the movement onset, motor-related regions (Mdl, Mdm, PMdm, PMdl, Sdl) showed stronger differences between baseline and action in the left hemisphere compared to the right hemisphere, which is an expected result as the subject performed the task with the right hand. In the occipital lobe, highest values of MI were reached approximately 250ms before movement onset.

**Figure 5.**
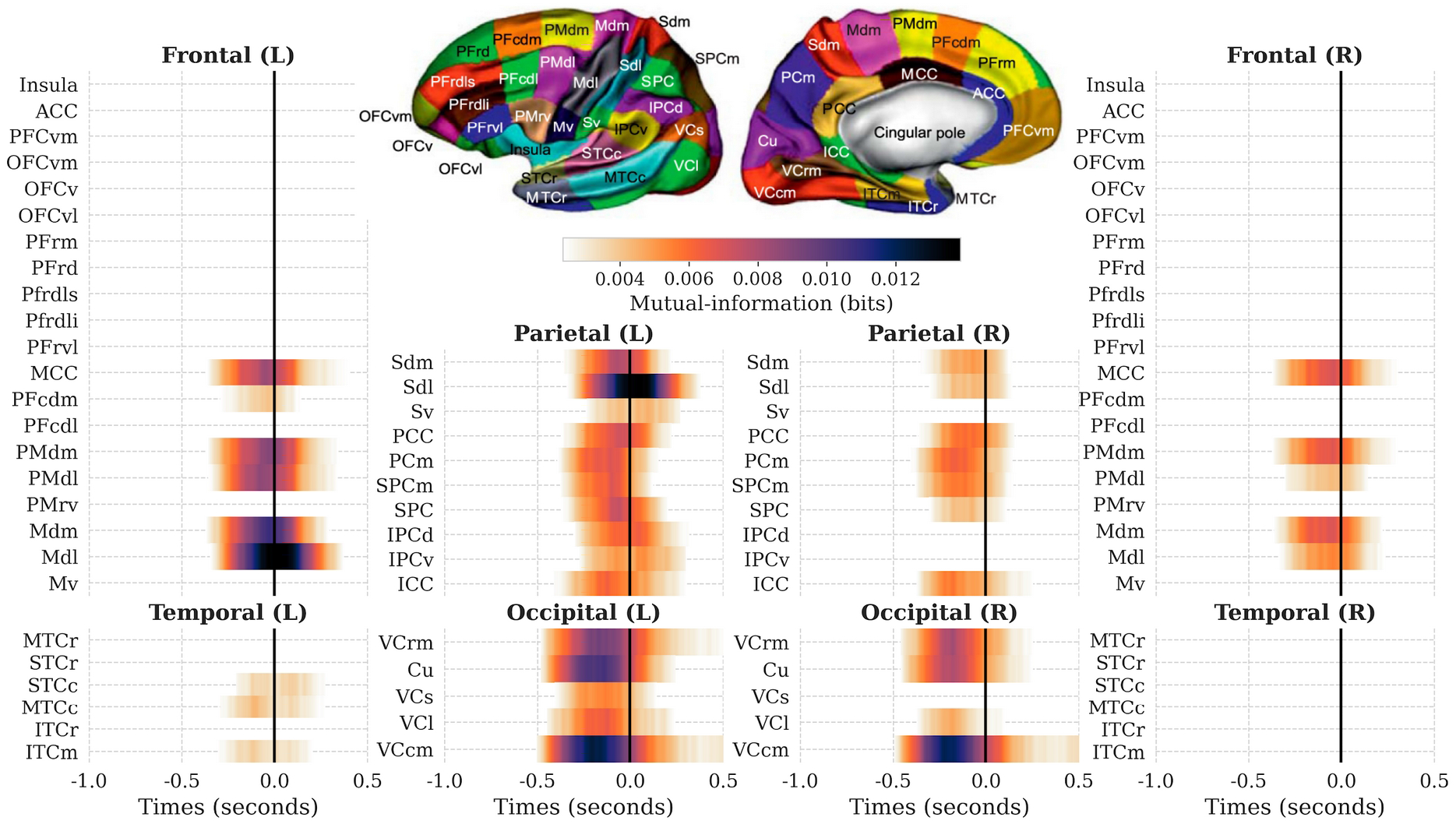
Brain areas showing significant differences of high-gamma activity between the action and the baseline. The plotted mutual-information is obtained by taking the mean across single-subject MI estimations, at each time-point and parcel. P-values are corrected for multiple comparisons using a cluster-based correction and displayed temporal clusters are inferred by thresholding the p-values at a significant level of p<0.05. Data are aligned on movement onset represented here with a vertical black line at 0 seconds.

To illustrate the proposed group-level statistical method on non-information-theoretic measures, we applied it on machine-learning (ML) outputs where a new classifier was systematically redefined and retrained at each time point and brain region for each subject (Sec. 2.5.1). To this end, we performed the same analysis as above (classifying action versus baseline in somatosensory (Sdl), motor (Mdl) and visual (VCcm) parcels, all three located in the left hemisphere (**Figure 6**). Similarly to the MI approach, permutations were computed by training and testing the classifier (LDA) on a randomly shuffled version of the label vector (Combrisson and Jerbi, 2015). Finally, we applied the random-effect group-level analysis on the AUC to infer p-values. Group-level statistics on both MI and ML scores lead to very similar significant temporal clusters with the only exception of VCcm, where the significant cluster identified by ML was narrower.

**Figure 6.**
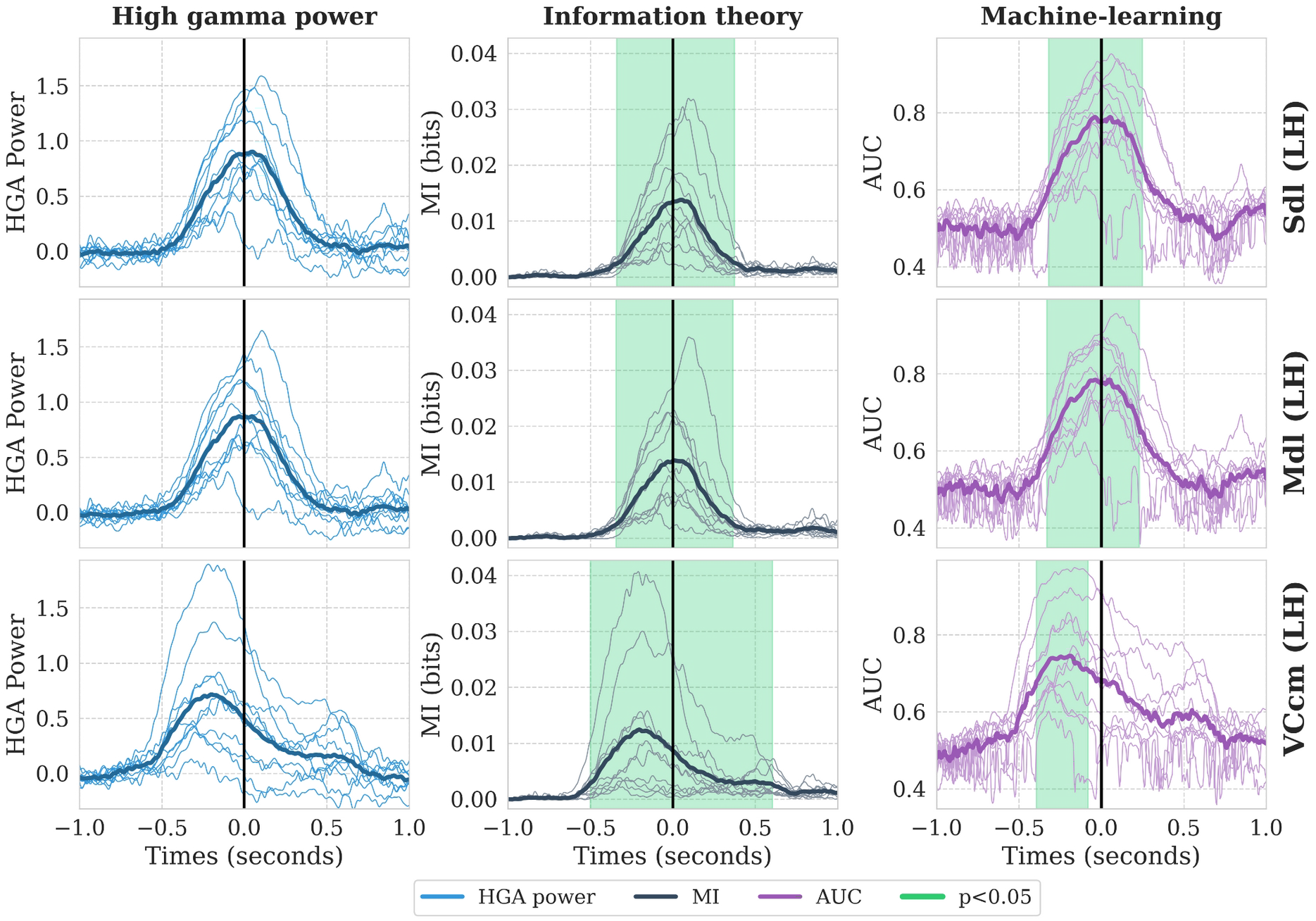
highlighting significant differences of high-gamma activity between the baseline and the action using group-level analysis on information theoretic and machine-learning metrics for three brain areas (dorsolateral somatosensory and motor cortex (Sdl, Mdl) and caudal medial visual cortex (VCcm)). The columns respectively refer to the high-gamma activity (blue), the mutual-information (black) and the machine-learning results (purple) expressed using the area under the curve (AUC). For all columns, thin lines stand for single-subject estimations and thick lines represent the mean across subjects. Green clusters highlight significant differences at p<0.05. Data are aligned on movement onset represented here with a vertical black line at 0 seconds.

We then tried to replicate previous visuomotor-related undirected and dynamic (time-resolved) functional connectivity (DFC) analyses (Figure 4 in Brovelli et al., 2017). We tested the hypothesis that interareal DFC was different during the execution of visuomotor associations with respect to baseline. Visuomotor-related changes in DFC with respect to baseline were estimated by using MI between the DFC and the discrete vector specifying if the trials were coming from the baseline or action condition. We used the RFX group-level approach, with p-values corrected for multiple comparisons across time and pairs of brain regions using the FDR correction. The time-averaged DFC for exhibited significant effects over the fronto-parietal network, mainly within the left hemisphere, with a strong implication of motor and premotor regions, superior parietal cortex such as strong bilateral connections within the occipital lobe (**Figure 7A**). We replicated the temporal evolution of the connectivity strength by computing the mean over pairs of brain regions exhibiting a significant contrast with the baseline (**Figure 7B**). In addition, the time-course averaged across pairs displayed a first peak occurring slightly after −0.4s, during the movement planning phase after stimulus presentation, a dissolution of the network around 0s and finally the emergence of a motor-related network at 0.2s when subjects performed the finger movement. In addition to the original paper, we computed the number of significant links according to the group-level model (FFX and RFX), such as approaches for correcting for multiple comparisons (FDR, Maxstat and Cluster-based across time points) (**Figure 7C**). Overall, the number of significant links was much larger when using the FFX model. Similarly, FDR correction led to a much larger number of significant edges, followed by the maximum statistics and finally the cluster-based correction.

**Figure 7.**
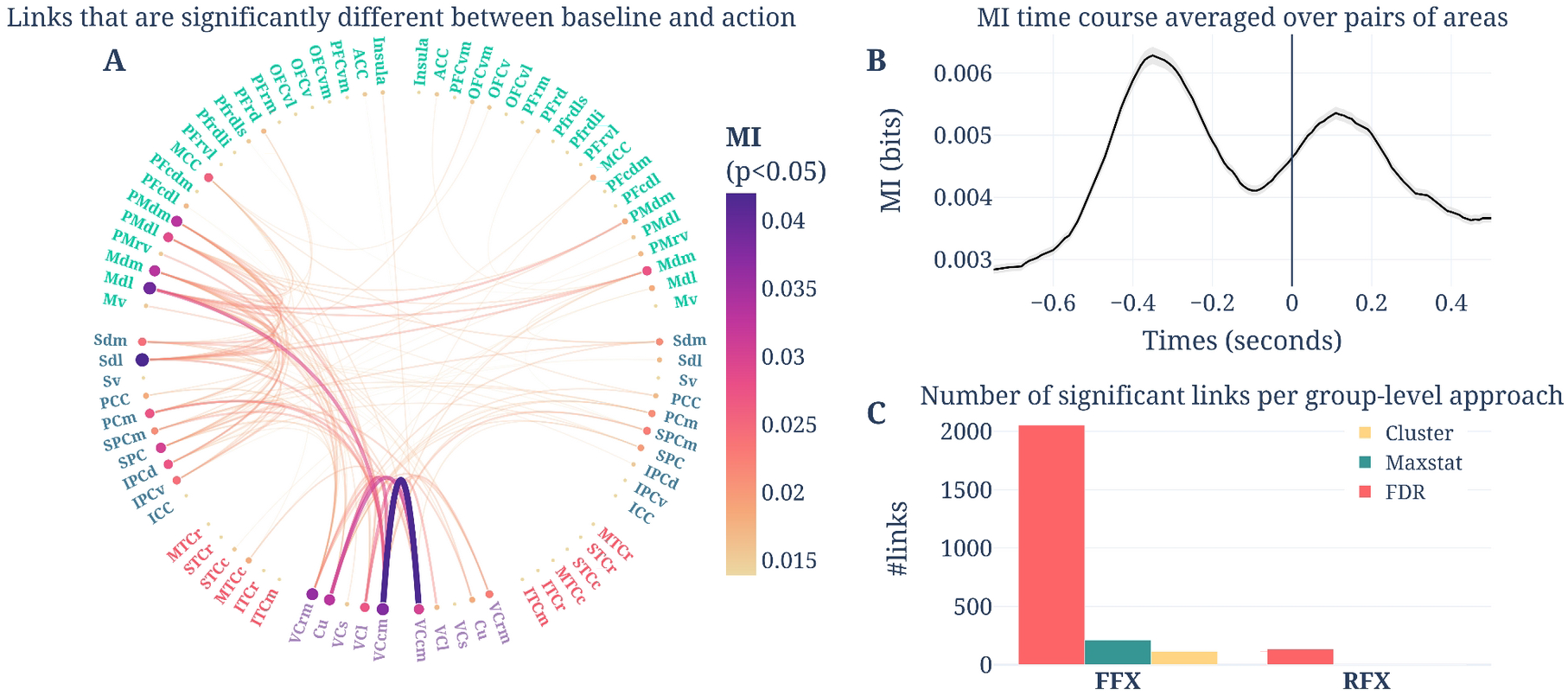
significant task-related undirected pairwise connectivity using MI (*A*) circular representation of the connectivity matrix of mean MI over time points (*B*) time-course of mean MI and 95% confidence interval across pairs of brain regions. Data are aligned on movement onset represented here with a vertical black line at 0 seconds and the time vector represents the center of each 500ms sliding window. (*C*) number of remaining significant links for the FFX and RFX models, respectively combined with FDR, maximum statistics and cluster-based corrections. All subplots were obtained after selecting the significant pairs of brain regions (p<0.05, corrected)

Finally, we investigated task-related network properties using standard graph-theoretical metrics (Sec. 2.5.1). To this end, we estimated the single-trial network efficiency, modularity and assortativity during both action and baseline periods. As for the MI timecourse of **Figure 7B**, all of the three metrics led to two peaks. We found a decrease of network efficiency (**Figure 8A**) and an increase both in network modularity and assortativity (**Figure 8B-C**) with respect to baseline period. We then computed the MI between the singletrial graph measures and a vector of labels indicating whether each trial was coming from the baseline or action periods. For the model of the group, we used a random-effect with cluster-based correction for multiple testings. For each of the three MI time courses, we found two significant clusters prior and after movement onset reflecting significant task-related differences of efficiency, modularity and assortativity.

**Figure 8.**
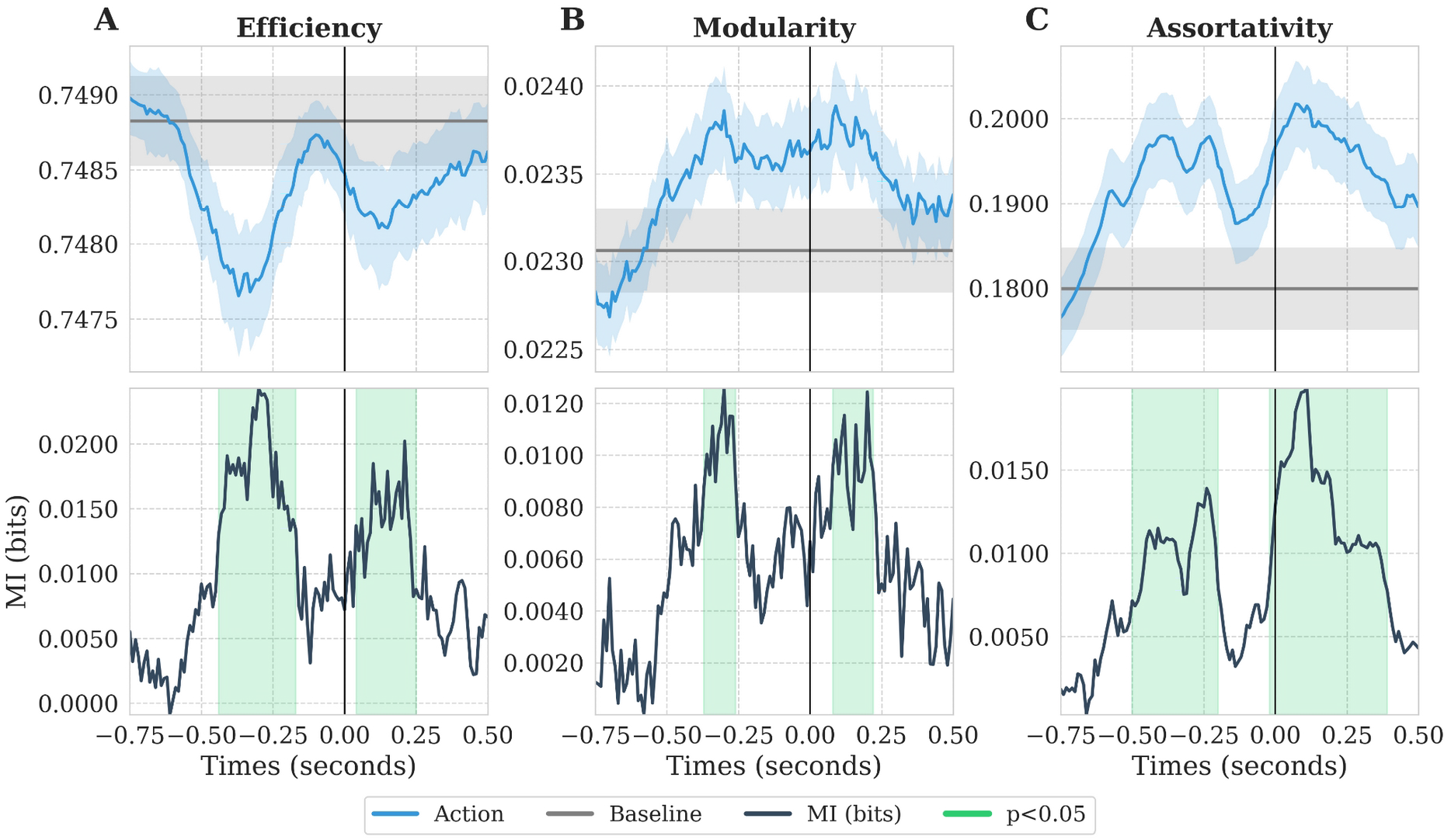
significant task-related graph metrics respectively measuring *(A)* the network efficiency, *(B)* the network modularity and *(C)* the network assortativity. Top row blue and gray lines represent the time courses of the graph measures respectively estimated during action and baseline. Shaded areas represent the 95% confidence interval estimated across subjects. Bottom row contains the MI time course and the significant clusters are highlighted in green (RFX - p<0.05). Data are aligned on movement onset represented here with a vertical black line at 0.

#### 3.2.2. Reinforcement learning system and intracranial EEG

We then tested our approach on intracranial HGA estimated from SEEG data recorded from the anterior Insula (aINS) in a reinforcement learning task. We computed the MI between the HGA power, estimated per bipolar contact, and the outcomes during both the reward (+1€ vs. +0€) and punishment (−0€ vs. −1€) conditions (**Figure 9**). As for the group-level statistics, we used a random-effect approach across contacts with cluster-based corrected p-values. Mean HGA power across subjects in the aINS showed clear modulations when receiving rewarding and punishing outcomes (+1€ and −1€) in contrast to +0€ and −0€ outcomes (**Figure 9A**). Overall, the difference of HGA power between outcomes was larger during the punishment condition which was then confirmed by the magnitude of MI and t-values (**Figure 9B-C**). The MI showed a significant peak occurring around 500ms after outcome presentation during both reward and punishment conditions. To further explore the consistency of the effect, we detected temporal clusters within-contacts and reported the proportion of contacts displaying a significant effect (**Figure 9D**). Interestingly, more than half of the contacts in the anterior insula showed significant differences between outcomes in the punishment condition compared to 20% during the reward.

**Figure 9.**
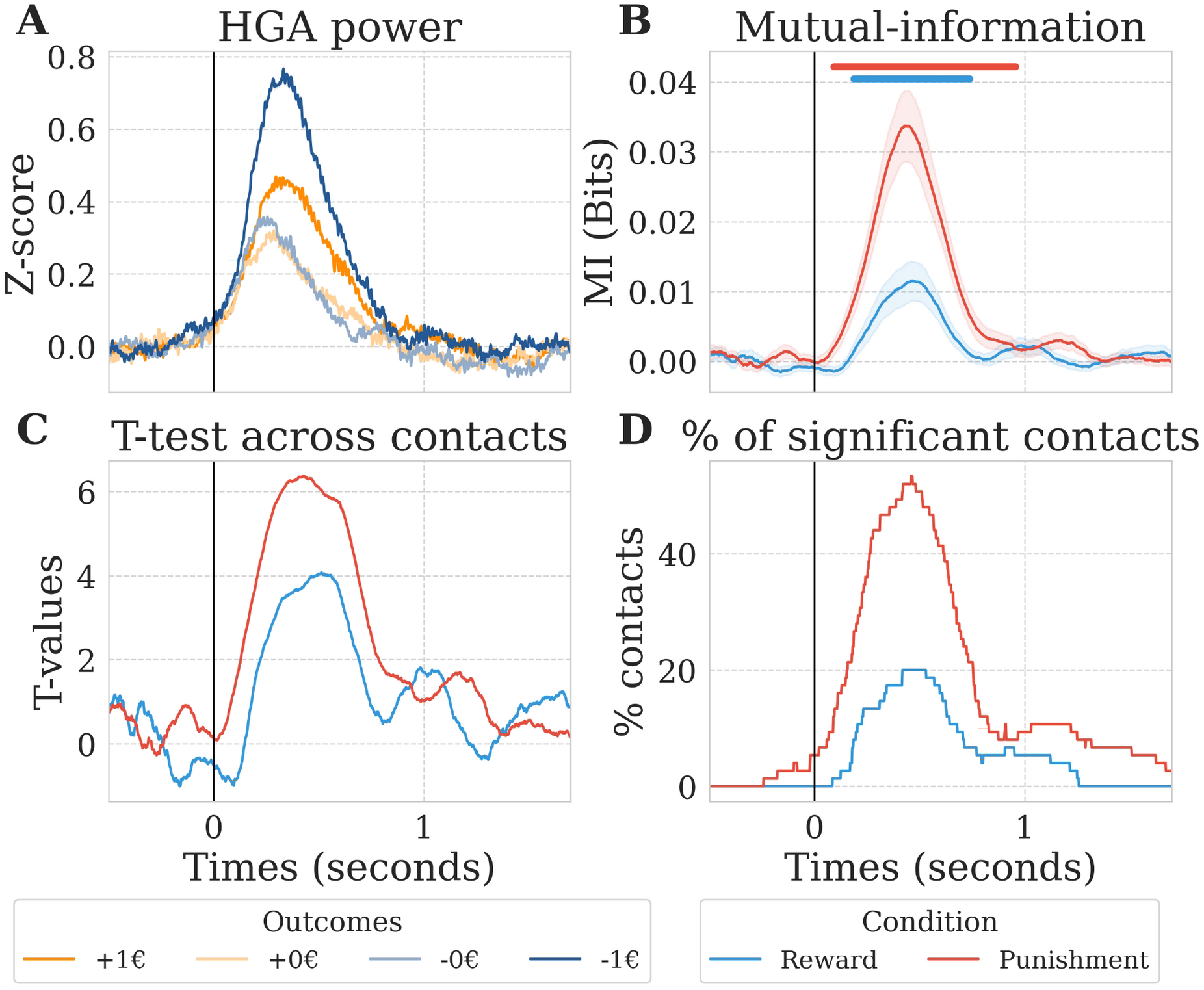
significant differences of HGA power between reward and punishment conditions in the anterior insula (13 subjects, 75 contacts). *(A)* Mean relative HGA power across subjects per outcome (+1€ and +0€ in dark and light orange, −1€ and −0€ in dark and light blue) (*B*) Mean MI computed between the HGA power and the outcomes during the reward condition {+0€, +1€} in blue and during the punishment condition {-1€, −0€} in red. Blue and red thick lines represent the significant temporal clusters obtained, at the group-level at p<0.05, respectively during reward and punishment conditions. (*C*), T-values obtained by computing a one-sample t-test contrasting single contact MI estimations against permutation mean (*D*) Proportion of contacts with significant effect. Vertical black line at 0 second represents the outcome presentation.

## 4) Discussion

In this study, we presented a statistical framework and associated toolbox called *Frites* for group-level inferences using non-negative information-based measures (e.g mutual-information, decoding) to identify cognitive brain networks. Our framework extends a previous approach for group-level inferences on information-based measures (Giordano et al., 2017) to encompass neurophysiological recordings with both homogeneous spatial samplings across participants (or sessions), such as whole-brain source level data M/EEG data and spatially sparse recordings, such as intracranial sEEG, ECoG and LFPs. In addition, it allows fixed- and random-effect models to adapt to inter-subject variability and uses non-parametric permutations with test- and cluster-wise correction for p-values inference. We used simulated data to compare the group-level approaches and complementary methods for correcting for multiple comparisons. We then tested our approach on human data to reproduce and extend existing results based on MEG data acquired during a visuomotor task (Brovelli et al., 2017, 2015) and spatially sparse intracranial data for investigating neural correlates of probabilistic learning (Gueguen et al., 2021). Finally, we illustrated how this framework can be applied at the level of local neural activity, on pairwise functional connectivity links or on measures summarizing network properties.

### 4.1 Comparison of group-level approaches

To compare F/RFX combined with different methods for correcting for multiple testings (maximum statistics, FDR and cluster-based), we defined three simulated groundtruths covering different scenarios : *(i)* weak and spatially diffuse effect; *(ii)* strong and spatially focal effect; *(iii)* varying spatio-temporal effect. For assessing the accuracy of the group-level approaches in retrieving the ground-truths, we used the Matthews Correlation Coefficient (MCC), which returns a single value reflecting the overall statistical performance by combining both sensitivity and specificity. Intuitively, the accuracy of both FFX and RFX increased with the number of subjects and the number of repetition per subject. For the same dataset (i.e., for the same number of subjects and repetitions), the FFX led to slightly higher accuracy than the RFX. However, this result should be nuanced by the fact that the first simulation used a highly reproducible effect across subjects. Indeed, when introducing inter-subject variability, FFX performance dropped until the effect started to be highly reproducible across subjects (~80% in the context of our simulations). This is explained by the fact that FFX is a single-level model, where the effect is estimated across subjects (Penny and Holmes, 2007) and cannot adapt to the inter-subject variability. On the other hand, RFX performance was stable at every level of inter-subject variability. Indeed, the RFX is a two-level model, in which the first level explains the effect size at the single-subject level and the second fits the population-level variance. Such capabilities to adapt to the variability and being able to generalize results to the population explain why, when possible, the RFX was preferred over the FFX (Lazar et al., 2002). While the RFX approach allows generalization to new subjects and accounts for random variations from subject to subject, it also comes at the cost of a lower sensitivity compared to the FFX which should be counterbalanced by a larger cohort.

Results from simulations also showed that methods for correcting for multiple comparisons were equally impacted with increasing sample-size or by the inter-subjects variability. However, maximum statistics and FDR were more impacted than the cluster-wise correction as the proportion of subjects having the effect was reduced. Note that while we focus here on inference on the population mean, which is the traditional focus in neuroimaging, the proportion of subjects showing an effect is also a valid target for statistical estimation at the population level (Ince et al 2021). In all scenarios, the overall performance of cluster-wise correction outperformed test-wise corrections. For the cluster-based correction (Bullmore et al., 1999; Maris and Oostenveld, 2007) the test is performed over the mass of the clusters, not on their precise spatiotemporal locations, therefore the p-value of the cluster is not representative of the member of the cluster. As a consequence, it was recently highlighted that numerous studies are reporting overestimated precisions of the statistical claims (Sassenhagen and Draschkow, 2019). To overcome this limitation and also to overcome the definition of the cluster forming threshold, several alternatives have been proposed like the Threshold Free Cluster Enhancement (TFCE) and probabilistic TFCE (Smith and Nichols, 2009; Spisák et al., 2019) or the non-parametric threshold free LISA framework (Lohmann et al., 2018). Overall, TFCE has been shown to provide a better sensitivity compared to standard cluster extent (Noble et al., 2019; Pernet et al., 2015). Future work could investigate how those modern cluster definitions perform once combined to the statistical framework proposed here.

### 4.2 Improving reproducibility through statistics

It has been shown that almost every study is unique in its analytical and statistical approach (Carp, 2012). This diversity is understandable considering the variety of designs and neurophysiological recordings. However, such diversity is a potential source for the lack of reproducibility (Gilmore et al., 2017; Puoliväli et al., 2020). The diversity in statistical approaches is further extended by the known differences between parametric and non-parametric statistical approaches. Data-driven non-parametric approaches are more likely to provide a better control for the FWER or FDR (Hayasaka, 2003; Nichols and Hayasaka, 2003; Puoliväli et al., 2020; Thirion et al., 2007). On the other side, it was also argued that parametric statistics can achieve similar sensitivity and specificity than nonparametric counterparts (Flandin and Friston, 2019). However, this is true at the condition that parametric assumptions are met, otherwise, violation of assumptions of the test can result in higher but also unpredictable FWER (Eklund et al., 2016). Parametric tests can also be more conservative, which means that many effects might be missed, pushing us to only report the tip of the iceberg (Noble et al., 2019). This is raising the question about our capacity to detect both strong localized effects along with weak and spatially diffuse effects (Cremers et al., 2017).

While there is a growing global consensus around the quality of control of nonparametric approaches, there are also classic criticisms encountered like the computational cost, the lack of reproducibility and the exchangeability hypothesis (Flandin and Friston, 2019). All three arguments are true, but should probably be nuanced. The computational cost is a technical problem that can be compensated using modern computers and programming libraries (multi-cores, multi-nodes, GPU, caching computations etc.). When living in the age where deep learning is applied on millions of images, it should be doable to perform 1000 up to 10000 permutations and get the results within a few minutes. Concerning the lack of reproducibility, it is true that by running the analysis several times the permutation scheme varies and therefore, some significant results might change. However, several reasons can explain this variability. First, the number of permutations is too low leading to a poor sampling of the null distribution. Then, if the result is still unstable for a larger number of permutations, it is questionable whether it is robust enough to consider being reproduced in the future. Finally, to avoid p-hacking (i.e. abusing of analysis techniques until nonsignificant results become significant (Head et al., 2015)) by changing the distribution scheme it is always possible to fix the random state of the machine, like this is done in many softwares, which leads to replicable permutations schemes (Pedregosa et al., 2011). The biggest danger of permutations probably lies in the *exchangeability* hypothesis (i.e., the variable we consider *exchangeable* in order to generate a null distribution of effects achievable by chance). For a simple contrast like *baseline vs. task*, where the null hypothesis is that there is no difference between the two conditions, the distribution of permutations should be constructed by permuting the labels. In other cases, choosing a permutation strategy appears sometimes as a subtle choice that requires deeper investigations like for machine-learning approaches (Combrisson and Jerbi, 2015; Valente et al., 2021). An incorrect permutation strategy (e.g., shuffling a variable that contains a structure like a temporal autocorrelation) can lead to a null distribution made of small effect size that are then going to be to easy to exceed by the true effect, leading to overconfident results (Brookshire, 2021; Liegeois et al., 2017). That being said, it is a problem the community is well aware of and much has already been written on how to choose the most appropriate permutation strategy (Maris and Oostenveld, 2007; Nichols and Holmes, 2002; Winkler et al., 2014).

Most studies have been shown to be underpowered which clearly limit our capacity to reproduce results across teams (Button et al., 2013; Ioannidis, 2005; Szucs and Ioannidis, 2017). Increasing both the number of subjects and within-subject data by data sharing is crucial for increasing the probability of detecting true effects. However, a standardization of group-level statistics and multiple testing, supporting both local and network level inferences, combined with powerful measures of information for minimizing assumptions on data distributions is another avenue to investigate toward higher reproducibility.

### 4.3 Group-level inferences on functional connectivity estimations

We showed that the proposed statistical framework could be used on pairwise FC to retrieve the task-related visuomotor network previously found using a linear mixed model on FC links estimated using correlation (Brovelli et al., 2017). Here, we reproduced the findings by entirely staying within the information-theoretical framework. We should nevertheless raise a note of caution concerning the interpretation of FC measures computed on noninvasive techniques, such as EEG and MEG, which may suffer limitations, among which volume conduction and leakage are potential confounds (Bastos and Schoffelen, 2016). Indeed, pairwise GCMI suffers the same limitations of standard correlation-based FC measures. In fact, the GCMI between two continuous variables is equivalent to a Spearman rank correlation, as pointed out by Ince et al. (2017). If transient common inputs affect pairs of regions on which FC measures are computed, the proposed pairwise GCMI analysis will reveal a significant effect. Potential solutions have been proposed in previous studies depending on the type of common input problem. If the common input signal can be recorded and it is included as a time series of the dataset, a potential solution is the use of multivariate, rather than pairwise, approaches to FC analysis. Partial correlation (Colclough et al., 2015) or conditional Granger causality (Ding et al., 2006; Wu et al. 2013) are examples of these techniques. On the other hand, if the common source input is not recorded and in the case of intrinsic spatial leakage, potential solutions such as orthogonalisation preprocessing steps (Hipp et al., Nature Neurosci 2012) or the use of FC measures that reduce zero-phase instantaneous coupling, such as the imaginary coherence (Nolte et al 2004), may attenuate such biases.

We also found that p-value correction at the edge-wise level can dramatically influence the proportion of significant links according to the statistical design (i.e F/RFX and the method for correcting MC). It was shown that the choice of the parcellation scheme, with a possibly large number of links, might lead to a high proportion of false negatives (Marek et al., 2020). To improve the capacity of detecting true task-related FC, several studies are using cluster-based approaches for networks (Baggio et al., 2018; Vinokur et al., 2015; Zalesky et al., 2010) or using predefined large-scale networks (Noble and Scheinost, 2020). In addition, it was recently shown that the number of significant links and statistical power were also highly dependent on the level of inference (i.e. at the edge, cluster or network level) (Noble et al., 2021). This is an important limitation to consider for future studies investigating task-related functional connectivity as only the tip of the iceberg is going to be reported. Those new approaches and new levels work on the same ideas that corrections for MC are going to be less stringent once informations are pooled and therefore, improve our capacity to detect true cognitive brain networks at the cost of being less precise when discussing the results (Sassenhagen and Draschkow, 2019). Those new developments should still be compatible with the statistical framework proposed here.

### 4.4. Information-based as a common language for cognitive brain networks

Measures of information are encapsulating the statistical dependency between brain data and an experimental variable. Here, for illustrating the statistical framework, we mainly used metrics from information theory by means of the Gaussian-Copula Mutual Information (GCMI) (Ince et al., 2017). GCMI has several non-negligeable advantages: *(i)* it can detects monotonic linear and non-linear relationships; *(ii)* it supports any combination of uni/multivariate of continuous/discrete variables, quantifying relationships on a common meaningful effect size scale; *(iii)* it contains a parametric bias-correction for unbalanced contrasts (i.e., when the number of data points is different); *(iv)* it is computationally fast. In addition, it also allows conditional MI (CMI) for removing the influence of a discrete or continuous variable. Typical use cases of CMI could be the estimation of the amount of information shared between the brain data and a behavioral variable, while conditioning of a third variable In addition, CMI is at the heart of the estimation of partial functional connectivity measures and covariance-based Granger Causality metrics. Indeed, Transfer Entropy (Schreiber, 2000), which is defined as the CMI between the past of the source and the present of the target conditioned by the past of the source, is mathematically equivalent to Granger causality for gaussian variables (Barnett et al., 2009). The monotonicity assumption for applying the GCMI between two univariate continuous time-series raises the question of the potential application to resting-state and long time-series. While this is still an open-question, it was recently shown that the GCMI and the weighted pairwise phase consistency outperform compared to other metrics for measuring the connectivity between brain and peripheral signals (Gross et al., 2021). This study used relatively long time-series, between 1 to 9 minutes and also showed that with longer time-series the mean distance between the connectivity measure and the surrogate distribution also increased. That said, those results also depend on the stationarity of the underlying connectivity. For an illustration of cases where the GCMI is not able to identify the relation between variables, see Fig 2. Ince et al 2017 and the Python implementation comparing several information-based estimators^7^. Overall, the GCMI is a versatile and general approach including metrics for local and network-level brain data analysis.

The framework presented here could additionally be combined with more powerful and feature-rich measures of information, such as encoding and decoding models for out-of-samples generalization, fully non-parametric kernel-based MI (Moon et al., 1995) and recent metrics from the Partial Information Framework (PID) (Ince, 2017; Wibral et al., 2017; Williams and Beer, 2010). Once combined with PID metrics, it becomes possible to investigate motivated questions like if there is redundant or synergistic information between a pair of brain regions about a variable of the task and whether this information is generalizable to the population.

Taken together, the framework presented here has the potential to investigate complex task-related modifications of local neural activity, but also more ambitious characterizations such as task-related long-range coactivations and feature-specific directed information flow (Bím et al., 2020). Linking brain networks to cognitive functions has been highlighted as one of the major challenges to overcome in the future (Bassett et al., 2020). Group-level statistics combined with rich measures of information represent one avenue to address this challenge toward the characterization of cognition.

### 4.5. Multi-scale analysis of cognitive brain networks

We showed how the statistical framework allows population generalization from the local scale of brain region to the large scale of brain-networks for characterizing both task-related functional connectivity and graph-theoretical properties. It further adapts to wholebrain parcel-based analysis of any modalities, from spatially uniform recordings like EEG and MEG to highly non-uniformly sampled invasive recordings like sEEG or ECoG. Outside of the scope shown here, the framework also has potential applications to other neurophysiological recordings like Utah arrays (Brochier et al., 2018), human and animal large-scale LFPs (Dotson et al., 2018, 2017), single- and multi-unit data and fMRI data.

### 4.6. Software implementation

Our approach has been implemented in an open-source Python package called *Frites*^8^ (*Framework for Information Theoretical analysis of Electrophysiological data and Statistics*) designed for inferring cognitive brain networks by means of measures of information. The toolbox, partially developed during Brainhack events (Gau et al., 2021), includes an implementation of the proposed group-level statistical pipeline, with fixed- and randomeffect models such as test- and cluster-wise corrections. By default, *Frites* is using a tensor-based implementation of the GCMI for faster computations. We recently added the possibility to use external functions for estimating information, like standard correlation, measures of distances, scikit-learn models (Pedregosa et al., 2011) or IDTxl non-parametric kernel-based measures of MI (Wollstadt et al., 2018). The toolbox also includes functions for estimating undirected and directed (covariance-based Granger Causality) single-trial functional connectivity. For interoperability, the main functions and classes of *Frites* are also supporting MNE-Python epochs-related objects. *Frites* is entirely written around a relatively recent package called *Xarray* (Hoyer and Hamman, 2017), which can be seen as a generalization of *Pandas* (McKinney, 2011) for labelled multidimensional arrays. One of the features of *Xarray* is that it allows returning and saving outputs with attributes. Therefore, each output of *Frites*’ function and classes also contains the inputs defined by the user such as the most relevant internal variables used for computations. The open-source accessibility of the package and the ability to track both internal and external variables, contribute to a reproducible science as it allows sharing self-contained files that include all of the parameters used for understanding and reproducing a result. From a programming perspective, we also provide online documentation^9^ with detailed function and input descriptions such as script and notebook examples. Code blocks are well commented and follow PEP8 guidelines for code readability. Finally, to improve long term sustainability as other recent Python packages (Combrisson et al., 2020, 2019; Meunier et al., 2020) we included both smoked (i.e. testing function’s input types) and functional unit tests through a continuous integration protocol (current coverage > 84%).

### 4.7 Limitations

The present work has several limitations and possible extensions that could be considered in future works. We used simple gaussian-based simulations to compare group-level approaches in terms of accuracy of ground-truth retrieving. Even if we simulated multiple scenarios of effect distribution, it is not straightforward that the presented results will behave the same in the context of real data. Future work could also investigate how this framework behaves in case of noisy sinusoidal time-series using auto-regressive models (Ding et al. 2006). The present work also investigated the number of required subjects, repetitions and proportion of subjects having the effect to achieve robust statistical inferences. Again, our simulation-based results illustrate a trend and can not be used to make sample size recommendations nor replace a proper power study (Baker et al., 2020; Maxwell et al., 2008). Future work could consider using more realistic data generation with a more sophisticated inter-subject variability modelisation. We also showed how the framework could be used on spatially non-uniform data like the sEEG. However, for both FFX and RFX inferences, measures of effect size could be unequally affected by differences in signal to noise (SNR) ratio. A possible workaround to investigate in the future could be to uniformize the SNR across brain regions by combining bootstrapping and stratification techniques by means of subsampling such that the distributional properties are made as similar as possible (Bosman et al., 2012).

## 5) Conclusion

There is a rising concern about our ability to reproduce results, partly because of underpowered studies and a wide range of analytical and statistical pipelines. Here, we proposed a statistical framework for group-level inferences on information-based measures of effect size. We demonstrated how the framework behaved according to various models of the group and corrections for multiple testing. We then illustrated the framework using information metrics coming from both the information-theory and machine-learning fields by reproducing and extending existing results on neurophysiological data. We also showed that the framework could be used at the level of brain regions, on functional connectivity and on measures of graph. The present work tends to provide more robust inferences on populations and therefore is a contributing piece toward more reproducible results.

## Abbreviations

FFX: Fixed Effect
RFX: Random Effect
MC: Multiple Comparisons
IT: Information-theory
MI: Mutual-information
ML: Machine-learning
MEG: Magnetoencephalography
sEEG: stereoelectroencephalography
FC: Functional Connectivity
DFC: Dynamic Functional Connectivity
GC: Granger causality
BIDS: Brain Imaging Data Structure
LFP: Local Field Potential
ECoG: Electrocorticography
FDR: False Discovery Rate

## Author contributions

Conceptualization : EC, BG, RAAI, AB

Data curation : EC, JB

Formal analysis : EC

Funding acquisition : AB

Investigation : EC, JB, AB

Methodology : EC, BG, RAAI, MA, AB

Project administration : EC, AB

Resources : AB

Software : EC, RB, RAAI, AB

Supervision : AB, BG, RAAI

Validation :

Visualization : EC

Roles/Writing - original draft : EC, BG, AB

Writing - review & editing : MA, RAAI, AB

## 6) Acknowledgments

EC and AB were supported by the PRC project “CausaL” (ANR-18-CE28-0016). This project/research has received funding from the European Union’s Horizon 2020 Framework Programme for Research and Innovation under the Specific Grant Agreement No. 945539 (Human Brain Project SGA3). MA, AB were supported by FLAG ERA II “Joint Transnational Call 2017” - HBP - Basic and Applied Research 2, Brainsynch-Hit (ANR-17-HBPR-0001). RB acknowledges support through a PhD Scholarship awarded by the Neuroschool. This work has received support from the French government under the Programme Investissements d’Avenir, Initiative d’Excellence d’Aix-Marseille Université via A*Midex (AMX-19-IET-004) and ANR (ANR-17-EURE-0029) funding. RAAI was supported by the Wellcome Trust [214120/Z/18/Z]. JB was supported by ANR-17-CE37-0018 and ANR-18-CE28-0016. Centre de Calcul Intensif d’Aix-Marseille is acknowledged for granting access to its high performance computing resources.

## Footnotes

1 https://brainets.github.io/frites/

2 https://brainets.github.io/frites/

3 https://brainets.github.io/frites/api/generated/frites.workflow.WfMi.html

4 https://brainets.github.io/frites/api/generated/frites.workflow.WfStats.html

5 https://brainets.github.io/frites/

6 https://brainets.github.io/frites/index.html

7 http://www.humanconnectomeproject.org/

8 https://brainets.github.io/frites/auto_examples/simulations/plot_ground_truth.html#sphxglr-auto-examples-simulations-plot-ground-truth-py

9 https://brainets.github.io/frites/auto_examples/estimators/plot_est_comparison.html#sphx-glr-auto-examples-estimators-plot-est-comparison-py

